# Epistasis in the receptor binding domain of contemporary H3N2 viruses that reverted to bind sialylated diLacNAc repeats

**DOI:** 10.1101/2024.11.26.625384

**Authors:** Ruonan Liang, Francesca Peccati, Niels L.D. Ponse, Elif Uslu, Geert-Jan Boons, Luca Unione, Robert P. de Vries

## Abstract

Since the introduction of H3N2 influenza A viruses in the human population, these viruses have continuously evolved to escape human immunity, with mutations occurring in and around the receptor binding site. This process, called antigenic drift, recently resulted in viruses that recognize elongated glycans that are not abundantly displayed in the human respiratory tract. Such receptor specificities hampered our ability to pick and propagate vaccine strains. Using ELISA, glycan array, tissue staining, flow cytometry, and hemagglutinin assays, this study revealed that the most recent H3N2 viruses have expanded receptor specificity by regaining effective recognition to shorter glycans. In recent H3 strains, Y159 and T160 are responsible for restricted binding to elongated glycans; in contemporary strains, however, Y159N and T160I dominate with a consequent loss of strength in receptor binding. Yet, effective receptor interaction is rescued by a remote mutation in the 190-helix, Y195F. The results demonstrate epistasis of critical residues in three of the four structural elements composing the HA receptor-binding site (the 130-loop, 150-loop, and 190-helix), which synergistically contribute to shape receptor binding specificity. Interestingly, a positive correlation exists between binding to an asymmetrical N-glycan containing an α2,6 sialylated tri-LacNAc arm and binding to human and ferret respiratory tract tissues. Together, these results elucidate the epistatic nature of receptor binding specificity during influenza A virus H3N2 evolution.

## Introduction

Influenza A viruses (IAV), having a reservoir of aquatic birds, occasionally cross the species barrier to infect humans, historically sparking pandemics. IAV can be divided into 19 subtypes based on the hemagglutinin (HA) and neuraminidase (NA) glycoproteins on the surface of IAV^1,2^. HA recognizes sialic acids on the host cell surface and fuses the virus’s membrane with that of the host, while NA promotes the release of new virions from infected cells. Among the HA and NA combinations, three of them, H1N1, H2N2, and H3N2, have been known to circulate in humans for decades. Notably, the introduction of IAV from avian species to humans coincides with a binding specificity switch from α2,3-linked (avian-type) to α2,6-linked (human-type) sialylated glycans^3,4^.

H3N2 viruses have a remarkable ability to escape host immunity and, as such, have infected the human population during the last six decades. During that time, the virus evaded neutralization by antibodies due to antigenic drift, driven by the accumulation of mutations in the most exposed globular head of HA. This part of the HA protein contains the receptor binding site (RBS) composed of four structural elements: the 130-loop, 150-loop, 190-helix, and 220-loop. Thus, antigenic changes in and around the receptor binding site (RBS) of HAs^5^ avoid antibody neutralization and impact the interactions with sialylated glycans and, eventually, compromise effective receptor binding^8–12^. Over the last decade, H3N2 viruses demonstrated altered receptor specificity, resulting in a robust selectivity for branched glycans containing at least three LacNAc repeats^12–15^. A LacNAc moiety is a disaccharide unit composed of a galactose β4-linked to an N-acetyl-glucosamine. In complex multi-antennary N-glycans, LacNAc units can form repeats of multiple lengths.

Despite the low abundance of LacNAc repeats containing glycans in the human and ferret upper respiratory tract^16,17^, at the beginning of the current century, H3N2 viruses incorporated mutations in the 150-loop that guarantee effective receptor binding to sialylated glycans containing at least 3 LacNAc repeats^18^. Employing high-resolution structural studies, we and others demonstrated that the aromatic side chain of the HA residue at position 159 was instrumental for the interaction with the innermost LacNAc repeat^19,20^. Yet, in 2021-2022, the dominating H3N2 strain in the northern hemisphere influenza had another set of changes at positions 158-160. This antigenic site B now contained NYT158-160NNI, which eliminated the aromatic side chain of residue 159 and, as expected, resulted in poor binding to α2,6-linked sialosides in a glycan array format but maintained some binding, specifically to extended branched sialosides^15^. However, that study did not assess the other HA substitutions in this antigenic drift period, which likely compensated for the weak binding to preserve viral infection.

Since January 2020, specific mutations in the 190-helix (G186D and D190N) have coevolved in human H3N2 HAs. A recent contribution from the group of Nicholas Wu demonstrated that the epistasis between G186D and D190N maintains binding to α2,6 sialylated glycans and alters HA antigenicity^21^. Contemporary strains favor extended glycan receptors, with significant binding to sialylated di-LacNAc structures^20^, regaining hemagglutination properties critical for antigenic surveillance^22^. It is, therefore, crucial to understand, at the molecular level, how glycan binding specificity evolves in contemporary H3N2 influenza viruses.

In this study, we interrogated which HA residues are responsible for the glycan binding specificity changes during H3N2 evolution in the last decade. We first focused on 3C.3a and 3C.2a1 viruses, the latter split into the 3C.2a.1b.2a.2a.3a.1 clade, revealing the reversion to di-LacNAc specificity^22^. All current H3N2 viruses contain Y159N/T160I in their HA, which abrogates receptor binding. This diminished receptor binding was rescued by introducing Y195F, present in all currently circulating strains. We demonstrate strong epistasis between amino acids T131K, G186D, D190N, F193S, and Y195F to maintain binding to an asymmetrical α2,6 linked sialic acid tri-LacNAc containing N-glycan. This, and perhaps similar structures, appear to be displayed on human and ferret respiratory tract tissues, whereas the symmetrical version is not.

## Results

### Y159 and T160 mutants are responsible for glycan binding changes during evolution

We and others have recently shown the importance of the Y159 position, as it directly interacts with extended LacNAc structures^12,19,20^. This data is derived from glycan arrays and STD NMR approaches; more standard affinity measurements, such as direct ELISAs, are lacking. This lack is mainly due to the scarcity of N-glycan material; only minute amounts are necessary for glycan array printing and are, therefore, hardly available for other experimental approaches. Here, we selected several N-glycans and introduced biotin at the reducing end, enabling a straightforward streptavidin coupling in microtiter plates. All N-glycans terminate with human-type receptors; we generated symmetrical bi-antennary N-glycan with a mono-, di- and tri-LacNAc extension (**A, C, and E)** and the asymmetrical counters parts of **C** and **E** in which the α3 arm is extended with either a di-LacNAc (**B**) or tri-LacNAc motif (**D**). The A/Switzerland/9715293/2013 (A/CH/13) H3N2 virus is part of the vaccine during 2014-15 and was chosen as a representative of the 3C.3a clade; conversely, A/Singapore/INFIMH-16-0019/2016 (A/SG/16), part of the 2018-2019 Northern hemisphere flu vaccine, was chosen as a representative of the 3C.2a1 strain. A/CH/13 encodes amino acids NSK at position 158-160, and A/SG/16 encodes NYT at position 158-160 (Fig. 1A).

**Figure 1.**
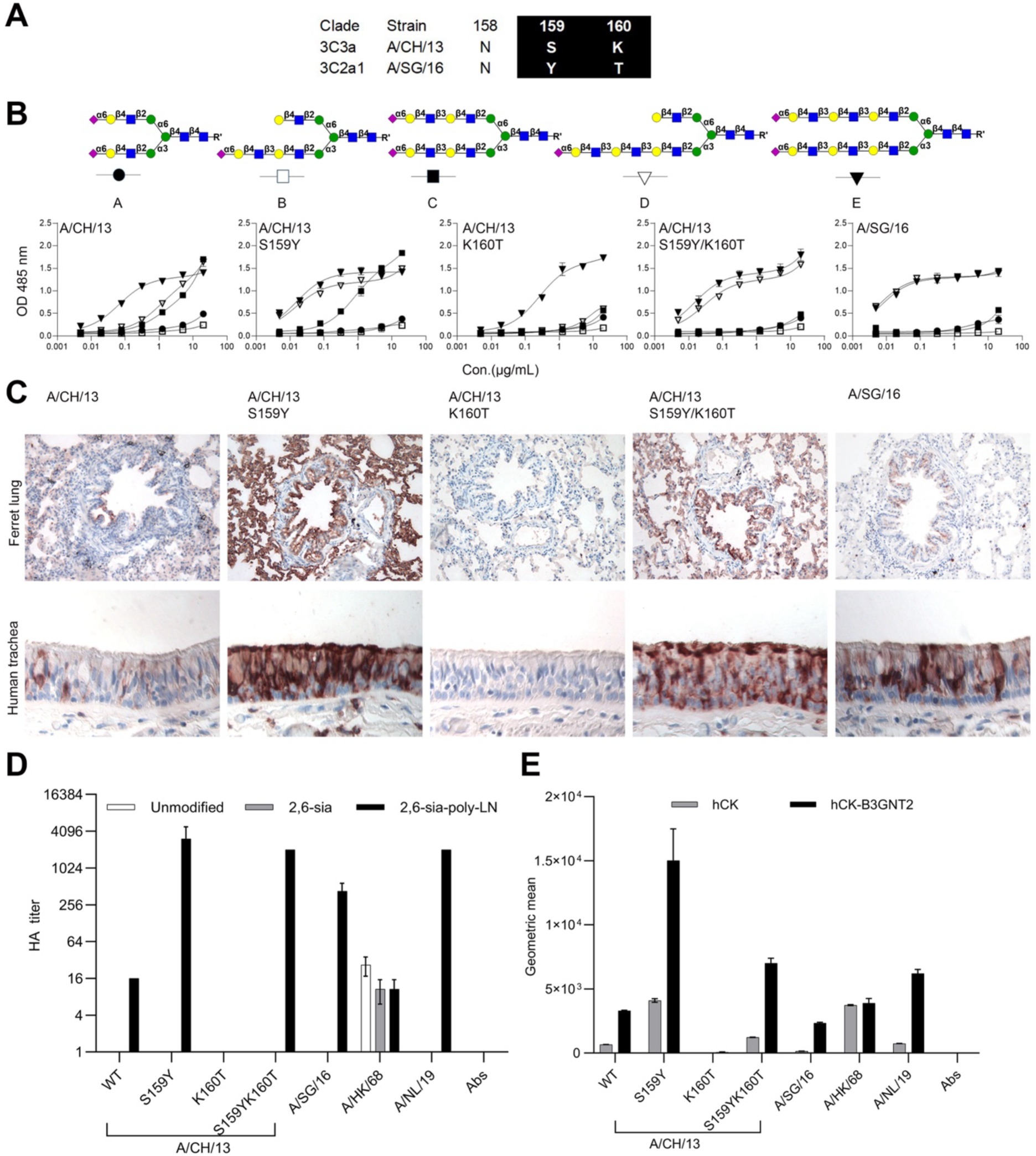
Y159 and T160 are responsible for glycan binding specificity towards LacNAc. **A**. Alignment of amino acids 159 and 160 between A/CH/13 and A/SG/16. **B**. Glycan structures A-E and binding avidities of A/CH/13 HAs (WT and mutations) and A/SG/16 to these structures was measured by ELISA. **C**. Tissue staining of A/CH/13 HAs (WT and mutations) and A/SG/16 WT to ferret lung and human trachea. **D**. Flow cytometry analysis of A/CH/13 HAs (WT and mutations) and A/SG/16 WT to hCK and hCK-B3GNT2 cells. **E**. Hemagglutination assay of A/CH/13 HAs (WT and mutations) and A/SG/16 WT to wildtype and glyco remodified chicken erythrocytes.

A/CH/13 wildtype HA preferentially binds the symmetrical tri-LacNAc compound **E** with less avidity to the symmetrical **C** and the asymmetrical tri-LacNAc structure **D** (Fig. 1B). A/CH/13 HA containing S159Y demonstrated an increased affinity to all compounds compared to the wild type, with the most notable increase in binding to **D**. However, when K160T was introduced, creating an N-linked glycosylation site, we observed a stark decrease in binding, and only compound **E** was bound. Reconstitution of S159Y/ K160T based on the A/CH/13 HA backbone, which mimics A/SG/16, showed a similar binding to asymmetrical **D** and symmetrical N-glycan **E**. Interestingly whereas A/CH/13 wildtype HA prefers compound **E**, the S159Y/K160T mutant does not differentiate between the symmetrical or asymmetric N-glycan. Quality control was performed using biotinylated Sambucus Nigra Lectin (SNA), which binds to α2,6 linked sialic acid attached to the terminal galactose, and all compounds containing that structure were similarly bound (Fig. S1A). At the same time, a symmetrical N-glycan in which the tri-LacNAc arms was not sialylated and a linear tri-LacNAc compound containing α2,6 linked sialic acid (6’SLN_3_-L) were used as controls (Fig. S1C), which showed that only the A/CH/13 S159Y mutant and SNA can bound the linear structure 6’SLN_3_-L. In contrast, all other HA proteins failed to bind to this compound.

Ferret lung and human trachea are biologically relevant respiratory tissues to analyze human influenza A virus receptor binding properties^23–26^. A/CH/13 wildtype HA bound the whole ferret lung weakly, and the S159Y significantly increased binding. Conversely, the K160T mutant failed to bind the ferret lung and human tracheal tissue sections. This mutant efficiently bound compound **E** in the ELISA, demonstrating that this structure is extremely rare in tissues. The double mutant S159Y/K160T gained binding to both tissues with less intense signals than the S159Y mutant. The A/SG/16 wildtype binding signal is restricted to the ferret bronchi, and we hypothesize that this is a biological difference between 3C3a and 3C2a.1 viruses. This data indicates that introducing a N-glycosylation site is detrimental to receptor binding. At the same time, the CH-pi interaction that the 159Y confers improved binding avidity to LacNAc and, therefore, can bind complex asymmetrical N-glycan **D**.

“Humanized” Madin-darby canine kidney (HCK) cells solely display glycan structures with α2,6-linked sialic acid^27^, the key receptors of human IAVs. We and others recently created HCK-B3GNT2 cell lines by overexpressing the B3GNT2 enzyme to increase the number of LacNAc repeat units, having a more human-like glycan profile^28–30^. The binding ability of the A/CH/13 and A/SG/16 was analyzed for these two cell lines using flow cytometry (Fig. 1D). A/HK/68 HA was used to detect α2,6-linked sialic acid presented in wildtype cells as this protein can bind α2,6 linked sialic acid presented on N-glycans as a mono-LacNAc structure^29^. Conversely, we used A/NL/19 HA protein control as this protein solely recognizes **E**^28^. A/CH/13 S159Y exhibited at least three times more binding to HCK-B3GNT2 cells than hCK cells. K160T abrogated binding to HCK-B3GNT2 cells, consistent with the ELISA and tissue staining results. The S159Y/K160T double mutant showed less binding than the S159Y mutant comparable to the A/SG/16 wildtype protein, and both exclusively bound to HCK B3GNT2 cells.

The hemagglutinin assay is a typical method to monitor IAV receptor binding; however, during antigenic drift, 3C.2a1 viruses have lost the ability to bind turkey erythrocytes^12^. We created glycoengineered erythrocytes to display N-glycans with elongated arms terminating in α2,6-linked sialic acid (2,6-sia poly-LN). We also made red blood cells that we treated with neuraminidase, and after that, α2,6 resialylated; therefore, these cells do not contain elongated LacNAc repeats (2,6-sia)^12^. We used these glycoengineered red blood cells to examine the agglutination titers of the A/CH/13 and A/SG/16 HA wildtype proteins and their mutants (Fig. 1E). As a control, A/HK/68, agglutinated wildtype, 2,6-sia and 2,6-sia poly-LN erythrocytes, and A/NL/19 only efficiently agglutinated 2,6-sia poly-LN erythrocytes. A/CH/13 wildtype only agglutinated 2,6-sia poly-LN erythrocytes^12^, importantly, S159Y and S159Y/K160T strongly agglutinated the erythrocytes having extended sialylated LacNAc moieties (2,6-sia poly-LN erythrocytes), similarly as A/SG/16 wildtype.

### Position 159 and 160 are asparagine and isoleucine in currently circulated H3N2 viruses, severely affecting receptor binding

Currently circulating H3N2 viruses have 22 amino acid mutations in HA1 when we compare the A/Netherlands/10595/2024 sequence with that of A/SG/16 (complete alignment is shown in Fig. S2). Amongst other amino acid changes in and around the receptor binding site, the changes at positions Y159N and T160I are noteworthy. This results in the loss of the CH-pi interaction between 159Y and the innermost galactose^19^, while the T160I results in a loss of an N-glycosylation site; both these mutations impact receptor binding properties^15^.

To analyze the receptor binding avidity changes to biologically relevant N-glycans, we expressed several mutants based on A/SG/16 HA, including Y159N, T160I, and Y159N/T160I, and employed these proteins in our ELISA approaches. The ELISA binding curves demonstrate that A/SG/16 Y159N abrogates binding to all the tested glycans compared to wild type, explaining the instrumental role of Y159 for efficient receptor binding (Fig. 2B). Instead, the T160I mutant exhibited a near-identical binding profile compared to the wild type, demonstrating that the presence or absence of N-glycan at that site does not interfere with receptor binding. Finally, the double mutant Y159N/T160I is bound efficiently to compound **E** and weakly to compound **C**.

**Figure 2.**
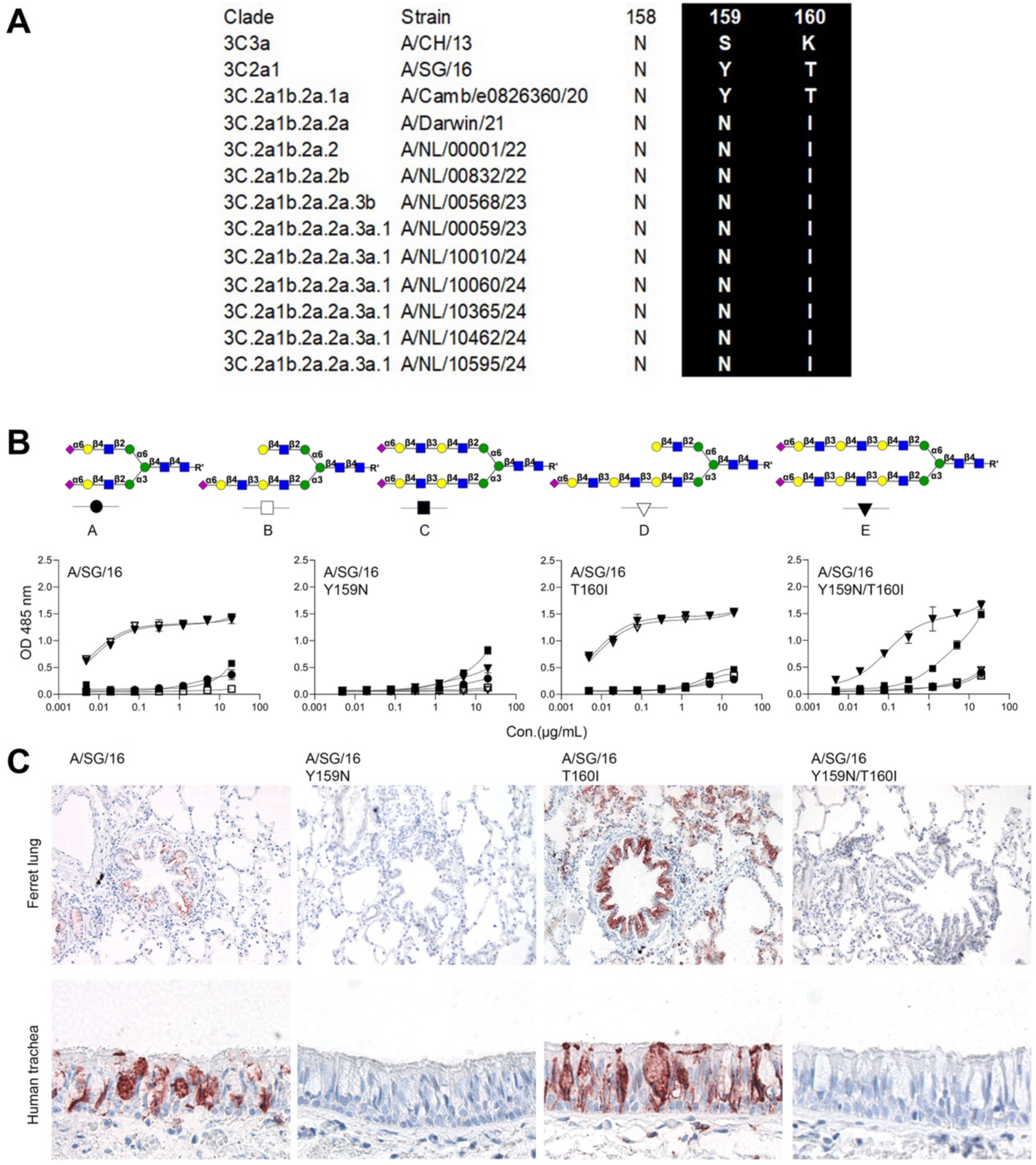
The binding specificities of A/SG/16 HAs (WT and mutations). **A**. Alignment of residues at 159 and 160 of 3C.2a.1 and its descendants. **B**. Binding avidities of A/SG/16 HAs (WT and mutations) measured by ELISA. **C**. The tissue staining of the A/SG/16 HAs (WT and mutations) to ferret lung and human trachea.

Next, we analyzed binding to ferret lung and human trachea tissue slides using the same A/SG/16 WT and mutant HA proteins. A/SG/16 WT bound to the bronchi in the ferret lung and goblet cells in the human trachea (Fig. 2C). The Y159N mutant failed to bind both tissues, as expected. Interestingly, the T160I binding profile significantly differed from WT as it could bind the alveoli in the ferret lung and bound in the epithelial layer of the human trachea. Y159N/T160I, on the other hand, completely failed to bind to both tissues while it did bind to compound **E**, indicating that this compound is hardly displayed in these tissues.

### F195 can rescue the binding profile of A/SG/16 Y159N/T160I

Next to Y159N and T160I, several other amino acid changes lie directly in the receptor binding site, which became fixed in currently circulating H3N2 viruses. These include T131K, G186D, D190N, F193S, and Y195F (Fig. 3A). It is unknown how T131K would affect receptor binding properties, but it is relatively conserved in human influenza viruses (45% of all H1, H2, and H3^31^). Amino acids at 186^21,32^, 190^5,33^, and 193^3,34,35^, on the other hand, are responsible for binding to LacNAc moieties in H3 and other subtypes. Finally, the Y195 is extremely conserved throughout all IAV subtypes and is considered part of the base of the receptor binding site with Y95, W153, H183, and L194^36^. To analyze their effect on receptor binding in contemporary H3N2 HA, we made single-point mutants (T131K, G186D, D190N, F193S, Y195F) in A/SG/16 and in the A/SG/16 Y159N/T160I background. The ELISA results demonstrate that T131K, F193S, and Y195F bound to compounds **D** and **E**. G186D exhibited weak binding to **E,** and D190N failed to bind at all (Fig. 3B). When the mutants were based on Y159N/T160I; the G186D, D190N, and F193S mutants lost binding to all compounds and the Y159N/T160I/T131K mutant bound with less avidity compared to the A/SG/16 T131K mutant. Surprisingly, Y159N/T160I/Y195F bound with very high avidity to compounds **C-E**; thus, the abrogation of binding by A/SG/16 due to Y159N/T160I is rescued by Y195F.

**Figure 3.**
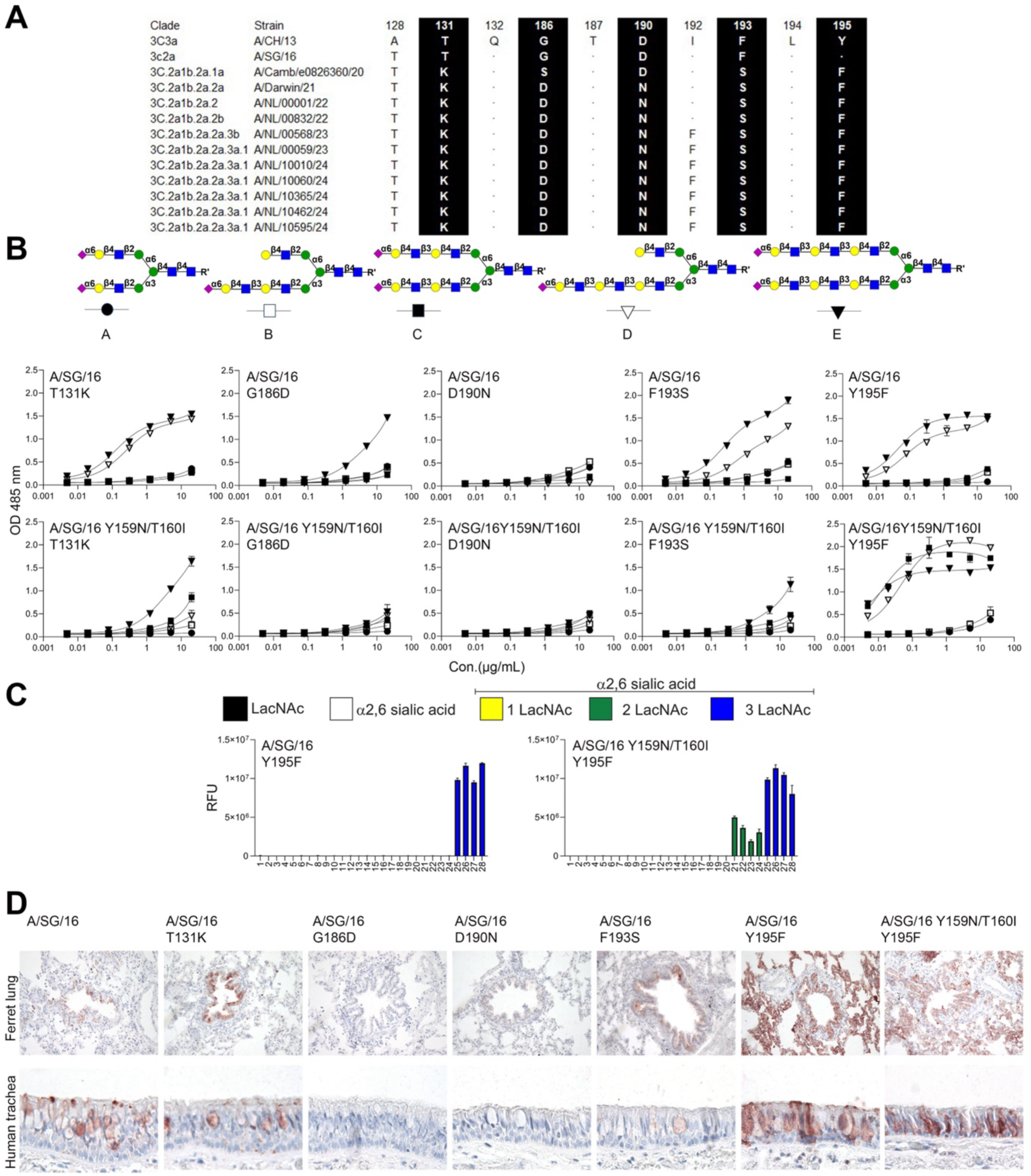
Y195F rescues abrogated binding due to Y159N/T160I. **A**. Alignment of the residues in 130 loop and 190 helix of H3N2 3C2a.1 viruses. **B**. Binding avidities of A/SG/16 and A/SG/16 Y159N/T160I mutant H3 proteins. **C**. Glycan array analysis of A/SG/16 Y195F and Y159N/T160I/Y195F. **D**. Tissue staining of A/SG/16 HAs (WT and mutations).

To further characterize the receptor binding specificities converted by the Y195F, we employed our previously published glycan array, populated with a broader variety of asymmetric N-glycans (Fig. S3)^22^. We demonstrate that Y195F without Y159N/T160I is only able to bind N-glycans carrying tri-LacNAc motifs and that only in the background of Y159N/T160I gains binding to N-glycans displaying di-LacNAc structures (Fig. 3C). Thus, the gain of binding avidity by Y195F only occurs when the 150-loop contains N159 and I160.

Additional flow cytometry analyses using hCK-B3GNT2 cells demonstrate that single mutants T131K, F193S, and Y195F are biologically active, while G186D and D190N are not (Fig. S4). In the background of Y159N/T160I, only the Y195F can bind hCK-B3GNT2 cells, further confirming the unique gain of binding afforded by Y195F.

To connect binding compounds **D** and **E** to binding to ferret lung and human tracheal cells, we conducted a tissue staining experiment using a selection of mutants: T131K, G186D, D190N, F193S, Y195F, and Y159N/T160I Y195F (Fig. 3D). A/SG/16 wildtype, T131K, and F193S exhibited the binding in the bronchi of the ferret lung and human trachea, whereas the G186D and D190N mutants did not. Y195F and Y159NT160I Y195F strongly bind in the whole ferret lung and human trachea. Conclusively, Y195F rescues the detrimental effect of Y159N/T160I on receptor binding.

### A case of exceptional epistasis between the 130, 150-loop, and 190-helix

To examine the functional interactions among the mutations, we constructed different constellations and assessed their binding activities. We created double mutations G186D/D190N and F193S/Y195F with or without Y159N/T160I, and assembled tri-, tetra-, and penta-mutations in the Y159N/T160I background. The ELISA analysis demonstrated that Y159N/T160I did not affect the abrogation of binding due to G186D/D190N, as hardly any binding was observed (Fig. 4A). F193S/Y195F and Y159N/T160I/F193S/Y195F showed strong binding to compounds **D** and **E**; the latter could also bind to **C**. The Y159N/T160I/D190N/F193S/Y195F mutant weakly bound to **E**, and binding was hardly improved with the introduction of G186D, perhaps slightly to compound **D**. With the additional introduction of T131K, resulting in Y159N/T160I/T131K/G186D/D190N//F193S/Y195F (7-mutant) displayed strong binding avidity to compounds **C-E**. Since the World Health Organization recommended that vaccines for use in the 2023-2024 northern hemisphere influenza season include the A/Darwin/9/2021 (H3N2) strain, we used it as a control of the dominating 3C.2a.1b.2a.2a.3a.1 strain. The binding activity of A/Darwin/9/21 was almost identical to that of the 7-mutant.

**Figure 4.**
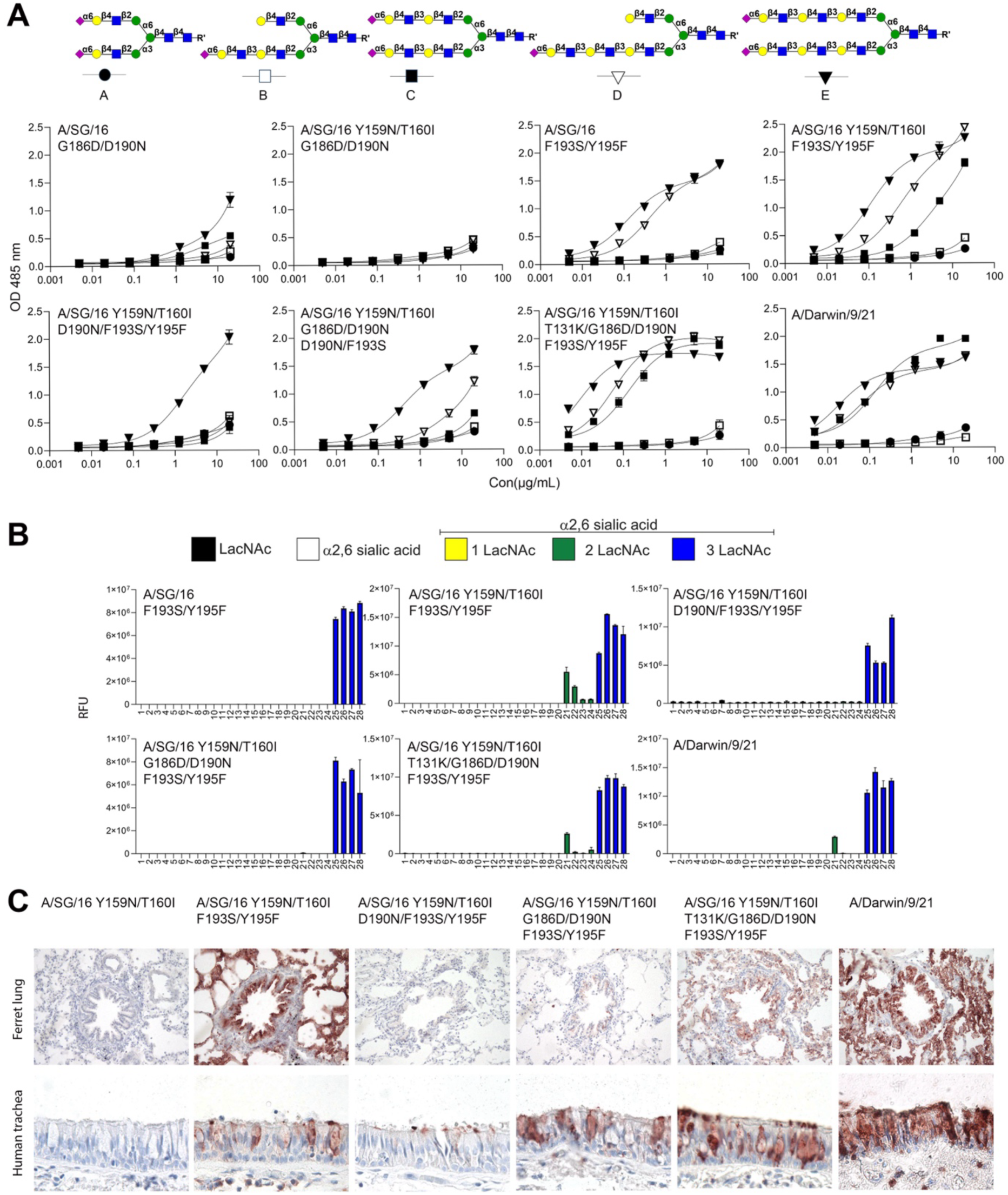
Epistasis between residues 131 186 190 193 and 195. **A**. Binding avidities of different mutational combinations. **B**. Glycan array analysis of H3 A/SG16 Y195F mutants. **C**. Tissue staining of the same HA proteins.

We wondered which of these mutants would convert binding to N-glycans with di-LacNAc motifs, as none of the proteins tested recognizes compound **B**. We thus performed glycan array analysis in which we demonstrated the F193S/Y195F only bound to N-glycans with tri-LacNAc structures. While the Y195F mutant in the background of Y159N/T160I confers binding to N-glycans with di-LacNAc motifs (Fig. 4B). Adding D190N (Y159N/T160I/D190N/F193S/Y195F) into this background resulted in a loss of di-LacNAc binding which was not regained by adding G186D (Y159N/T160I/G186D D190N/F193S/Y195F). As with the ELISA data, the additional introduction of T131K (Y159N/T160I/T131K/G186D/D190N/F193S/Y195F) exerted binding to a di-LacNAc motif, and binding was nearly identical to that of A/Darwin/9/21. This means that none of these mutations are indispensable. Remarkably, compound **21** was preferred over other di-LacNAc-containing structures. This difference to structures **22-24** is the additional introduction of LacNAc on the α6 arm (Fig. S3), probably creating steric hindrance.

Finally, we tested the mutants for binding to ferret lung and human trachea tissues. F193S/Y195F convert binding in the Y159N/T160I background, but D190N/F193S/Y195F and G186D/D190N/F193S/Y195F hardly or weakly bound (Fig. 4C). The latter, interestingly, only demonstrated binding to human tracheal tissues, which probably correlates to the ability of this protein to bind to **D** and some N-glycans with a di-LacNAc arm (**#21**). When T131K was introduced, the resulting T131K/G186D/D190N/F193S/Y195F mutant bound with high responsiveness to ferret lung and human trachea tissues, nearly identical to A/Darwin/9/21. The supporting data demonstrated that no proteins tested bound unsialylated or a linear tri-LacNAc compound (Fig. S5A). Flow cytometry analysis using both hCK and hCK-B3GNT2 cells revealed an overall preference for hCK-B3GNT2 cells in which we confirmed the epistatic interaction between Y159N/T160I and G186D/D190N, which are detrimental for receptor binding with T131K and Y195F rescuing binding. These seven mutants have nearly identical receptor binding properties to A/Darwin/9/21 (Fig. S5B).

### The structural ramifications of the Y159N/T160I and Y195F mutation revealed by molecular modeling

Y159N and T160I have apparent structural effects on receptor binding, including the loss of CH-pi interaction with the innermost galactose of tri-LacNAc-containing glycans and the loss of a glycosylation site. Y195 is exceptionally conserved in all IAVs and forms the base of the receptor binding site with Y98 and H183. To characterize how these mutations impact receptor binding at the molecular level, we performed all-atom microsecond molecular dynamics (MD) simulations of monomeric models of the HA RBS of A/SG/16 and A/SG/16 Y159N/T160I/Y195F bound to **D**. Fig. 5A shows an overlay of 120 MD frames illustrating the flexibility of the bound compound **D**. In A/SG/16, the CH-pi interaction between the aromatic ring of the Y159 and the innermost Gal engages the extended chain of **D** in close contact with the 150-loop. Instead, in the Y159N/T160I/Y195F mutant, the loss of CH-pi interaction, together with the lack of the glycosylation site at N158, destroys the intermolecular contacts between **D** and the 150-loop, suggesting a marginal role of this loop in binding **D**. Yet, at the sialic acid binding site, most of the HA-receptor interactions are preserved. Critical contacts for sialic acid binding include a hydrogen bond between O8 and the side chain of Y98 and a pair of hydrogen bonds between the sialic carboxylate group and the side chains of S136 and S137, located in the 130-loop (Fig. 5B). These contacts are preserved along the entirety of the simulations and show equivalent distance distributions in A/SG/16 and the Y159N/T160I/Y195F mutant.

**Figure 5.**
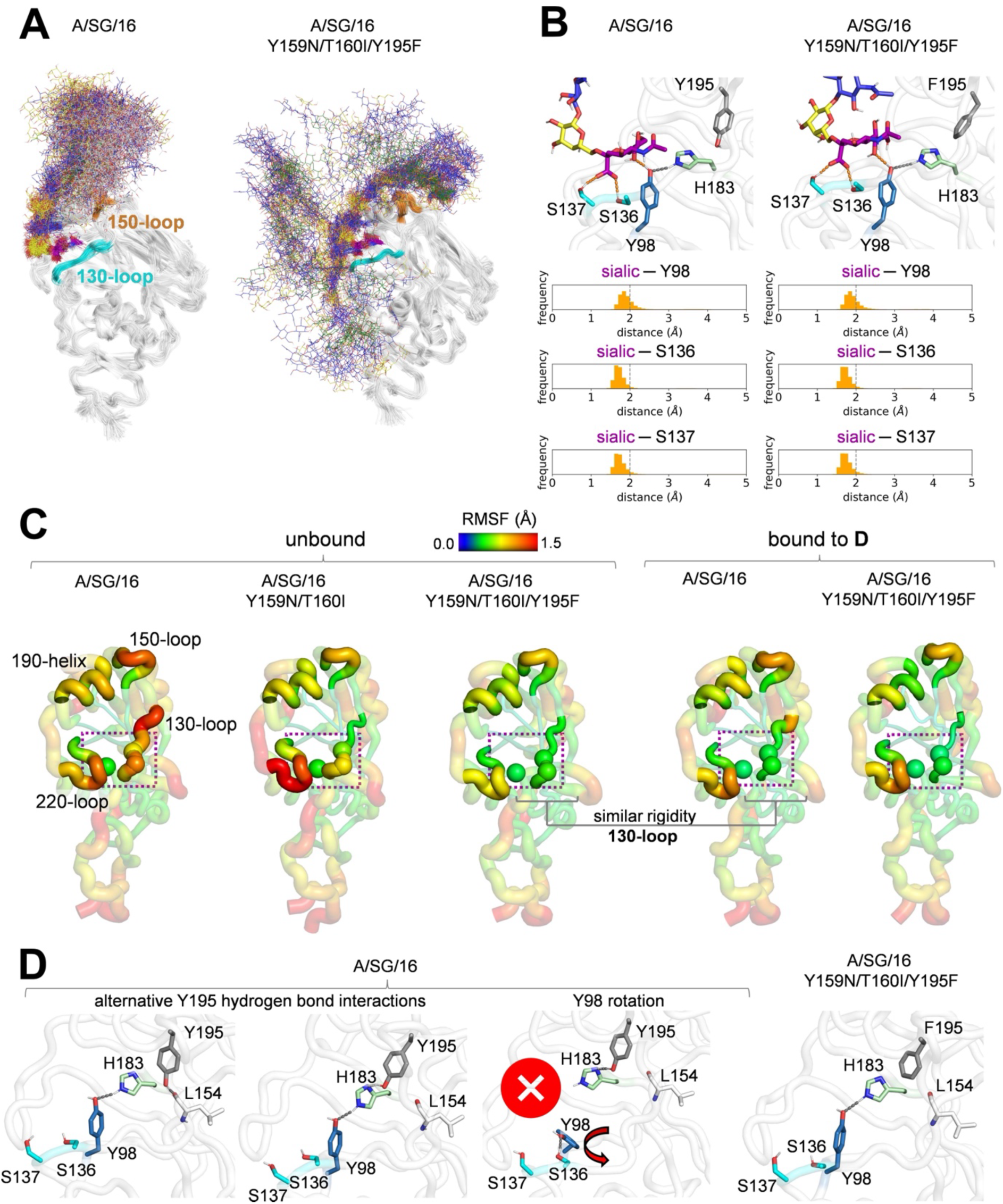
The Y195F mutation allosterically rescues A/SG/16 Y159N/T160I receptor binding. **A**. Overlay of 120 frames sampled with an even stride from three 400 ns MD replicas (1.2 μs total simulation time) of A/SG/16 and A/SG/16 Y159N/T160I/Y195F HA monomeric RBS models bound to compound **D**. **D** is shown as colored lines and the N158 glycan as white lines. The 130-loop and 150-loop are highlighted in cyan and orange, respectively. **B**. Distribution of hydrogen bond distances between the sialic acid of **D** and the side chains of Y98 (blue sticks) and of S136 and S137 (cyan sticks) along the same MD simulations. Relevant distances are shown as orange dashed lines. The side chains of H183 and Y/F195 are shown as green and grey sticks, respectively. **C**. Protein backbone root-mean-square fluctuations (RMSF) along the MD simulations (1.2 μs) of unbound HA monomeric RBS models of A/SG/16, A/SG/16 Y159N/T160I, A/SG/16 Y159N/T160I/Y195F, and the same models of A/SG/16 and A/SG/16 Y159N/T160I/Y195F bound to **D**. 130-loop, 150-loop, 190-helix, and 220-loop are highlighted. Cα atoms of Y98, S136, and S137 are shown as spheres. **D** and the N158 glycan are not shown for clarity. **D**. Selected frames along the MD simulations of unbound A/SG/16 and A/SG/16 Y159N/T160I/Y195F HA monomeric RBS models highlighting alternative hydrogen bond networks engaging Y98, H183, and Y195. Relevant residues are shown as sticks: Y98 in blue, S136 and S137 in cyan, L154 in white, H183 in green and Y/F195 in grey. The area involved in sialic acid binding is marked with a purple diamond.

We evaluated the flexibility of the four RBS elements (130-loop, 150-loop, 190-helix, and 220-loop) in unbound A/SG/16, Y159N/T160I, and Y159N/T160I/Y195F models and the models of A/SG/16 and Y159N/T160I/Y195F bound to **D** by computing backbone root-mean-square fluctuation values (RMSFs) along the MD simulations, shown in Fig. 5C. In the unbound state, the Y159N/T160I/Y195F mutant shows greater rigidity than A/SG/16 and Y159N/T160I in the 130-loop, and to a lesser extent in the 220-loop (the latter being also involved in sialic acid binding through S228). Additionally, the Y159N/T160I mutations enhance the rigidity of the 150-loop. RMSF values of residues S136 and S137 in the unbound Y159N/T160I/Y195F mutant are comparable to those of A/SG/16 and Y159N/T160I/Y195F bound to **D**, suggesting that the Y195F mutation compensates for the loss of affinity for the 150-loop by allosterically increasing the preorganization of the 130-loop, which is directly involved in sialic acid binding.

Finally, we analyzed the MD simulations of unbound A/SG/16 and Y159N/T160I/Y195F to interpret this allosteric effect at the atomic level. In A/SG/16, Y195 can alternatively engage in hydrogen bonding with either the backbone carbonyl of L154 at the base of the 150-loop or the Nδ of H183 (Fig. 5D). In turn, H183 forms a hydrogen bond with Y98, maintaining its side chain correctly oriented for sialic acid binding. Therefore, Y195 provides allosteric communication between the 150-loop and the region encompassing Y98 and the 130-loop, both involved in recognizing tri-LacNAc structures. The Y195F mutation breaks this long-range communication by ablating the hydrogen bond ability. Additionally, in 3% of the models in the A/SG/16 simulation, the Y98 side chain rotates away from H183, leading to a conformation that is not preorganized for sialic acid binding (Fig. 5D). Conversely, in the Y159N/T160I/Y195F mutant, Y98 remains preorganized for binding and at closer contact with H183 (≤ 4 Å) for the entirety of the simulation. Therefore, the ability of the Y159N/T160I/Y195F mutant to recognize both di- and tri-LacNAc structures is attributed to a higher affinity compared to A/SG/16 for the common sialic acid unit.

## Discussion

H3N2 viruses continuously evolve to escape population immunity, this process of antigenic drift occurs due to mutations in and around the receptor binding site^5,37^. Such mutations block dominant antibodies that neutralize the virus by inhibiting IAV sialic acid binding^38^. During decades of antigenic drift, human H3N2 viruses have acquired an exquisite receptor binding specificity for N-glycans carrying an α2,6-sialylated tri-LacNAc at the α3 arm^12,19^. During the last couple of years, however, these viruses reverted to bind di-LacNAc structures again, which coincides with hemagglutination properties, essential for antigenic surveillance^22^. Here, we determined the molecular determinants for this reversion of receptor binding and, to our surprise, found that a mutation of a highly conserved Y195 to an F resulted in di-LacNAc binding if the 150 loop contains N159 and I160. Conversely, positions 159 and 160 were vital in the receptor binding switch between 3C.3 and 3C.2a viruses and their descendants.

Residue 195 is exceptionally conserved in all subtypes, which made this substitution so intriguing in the 3C.2a.1b.2a.2a.3a.1. strains. Position 195 is part of antigenic site B and includes residues 159, 160, 186, 190, and 193, all independently affecting receptor binding. Residue 131, on the other hand, lies in antigenic site A and affects receptor binding. Single mutations at positions 186 and 190 that abrogate binding are epistatic, as previously shown^21^, while other single mutants hardly affect receptor binding. The only mutant able to rescue the binding of the Y159N/T160I change was Y195F. Emphasizing the importance of this conserved residue.

The residue at 194 is equally conserved and has been shown to undergo egg-adaptive mutations in vaccines, which are, therefore, less efficient^32,39,40^. The epistatic nature of positions 194, 186, and 190 has been demonstrated, as they are highly antigenic^21^. Interestingly, these HA proteins only bind N-glycans carrying an α2,6-sialylated tri-LacNAc at the α3 arm. Here, we demonstrate that in nature, these viruses revert to binding di-LacNAc structures, as these are presumably more abundant in the human upper respiratory tract^17^. Interestingly, while H3N2 antigenic drift forced the virus to bind low abundant complex N-glycans, these viruses did gain polymerase activity^41^. Thus, following naturally occurring mutations informs us of possible convergent and preferred receptor binding profiles. It has recently been shown that human H1N1 viruses evolved to bind near identical glycan structures^22,42^.

Our use of complex symmetrical and asymmetric N-glycans, displaying either two or three LacNAc repeats at the α3 arm in ELISA-based assays, is a first. We employed such structures because viruses equally bound compounds **D** and **E** in glycan arrays. Previously, it was postulated that the symmetrical structures should be preferred as they could engage in a bidentate manner^13^, however, with our recent studies, we clearly showed that the asymmetrical variant is often bound equally well. However, direct affinity differences have not been measured until now. Although our ELISA-based assays are still dependent on multivalency and thus will not resolve a Kd, we observed significant differences in binding avidity to these structures.

Interestingly, HA proteins that bound the asymmetrical compound **D** could also bind ferret and human respiratory tissues, while proteins restricted to the symmetrical structure **E** did not (Fig. S6). We thus like to hypothesize that both human H1N1 and H3N2 viruses preferentially evolved to bind di- and tri-LacNAc structures on the α3 arm, while the α6 is not extended, as it is these kinds of structures that are more abundant in the upper respiratory tract of humans.

These findings provide insights into the complex interplay of mutations affecting glycan binding during IAV evolution at the molecular level. Our data will aid in the functional and antigenic surveillance of ever-drifting H3N2 viruses and assist in picking future vaccine strains.

## Materials and methods

### HA expression and purification

HA encoding cDNAs, A/CH/13, A/SG/16, and A/Darwin/9/21 (synthesized and codon-optimized by GenScript) were cloned into pCD5 expression vector, with a signal sequence, GCN4 trimerization motif, a TEV cleavage site, the twin-strep (IBA, Germany) and a super folder GFP or mOrange2^43^. Mutations were introduced into HA genes by using site-directed mutagenesis. The HAs were expressed in the HEK 293S GnTI(-) by transfecting the expression vectors with polyethyleneimine I (PEI) (DNA:PEI is 1:8). After 6 hours of post-transfection, the transfection medium was moved and replaced by the 293 SFM II expression medium (Gibco), supplemented with 3.0 g/L Primatone (Kerry), 2.0 g/L glucose, 3.6 g/L bicarbonate, 0.4 g/L valproic acid, 1% glutaMAX (Gibco), and 1.5% DMSO. After 5-6 days, the Culture supernatants were collected. HAs were purified by the Strep-Tactin Sepharose beads (IBA, Germany) according to the manufacturer’s instruments.

### Enzyme-linked immunosorbent assay (ELISA)

Nunc Maxisorp 96 wells plates (Invitrogen) were coated with 50 μL of 5 μg/mL streptavidin (Westburg) in PBS overnight at 4°C followed by the blocking with 300 μL of 1% BSA in PBS-T for 3 hours at room temperature. The streptavidin-coated plates were coated with 50 μL of 50 nM biotinylated glycans overnight at 4°C, followed by the blocking with BSA again. HAs at 20 μg/ml were precomplexed with strepmab and goat-anti-human antibodies (Invitrogen (#31410)) in a 1:0.65:0.325 molar ratio on ice for 30 min. Proteins were added to the plates, diluted serially 1:3, and incubated for 2 hours at RT. HA binding was developed by OPD and stopped by the 2.5M H_2_SO_4_ after 5 min. The UV reader (Polar Star Omega, BMG Labtech) measured the optical density at 485 nm. The data was calculated from three independent experiments in duplicate.

### Glycan microarray binding of HA proteins

The glycan array microarray was performed as described previously^12,44^. HAs were precomplexed with human anti-streptag-HRP and goat anti-human-Alexa 555 antibodies in a 4:2:1 molar ratio in 40 μL of PBS-T on ice for 30 min. Subsequently, they were incubated on the glycan array surface in a humidified chamber for 90 min. Then, the slides were washed in PBS-T, PBS, and deionized water. After the water had been removed by centrifugation, the slides were immediately scanned. After removing the highest and lowest values, the data was calculated over four replicates.

### Immunohistochemistry

Immunohistochemistry was performed as previously described. Briefly, Ferret lung and human trachea sessions at 5 μm were deparaffinized and rehydrated, followed by an antigen retrieval step by heating the slides for 10 min in sodium citrate. The endogenous peroxidase was inactivated by 1% H2O2 in MEOH. Tissues were blocked by the 3% BSA in PBS overnight at 4°C. The precomplexed HA, strepmab, and goat-anti-human antibodies were incubated with tissues for 90 min at RT. For ferret lung, the HA was 50 μg/ml. For the Human trachea, the HA was 20 ug/ml. After incubation, the tissue was stained by AEC substrate for HRP (Abcam) and subsequently stained by hematoxylin.

### Hemagglutination assay

The chicken erythrocytes were remodified with different lengths of LacNAc, terminal α2,6 sialic acid described previously^12^. 20 μg/mL Hemagglutinin was incubated with strepmab and goat-anti-human HRP on ice for 30 min (same ratio as ELISA). Precomplexation was diluted serially in 2-fold.) 1% Erythrocytes were added and incubated at RT for 3-4 hours.

### Flow cytometry analysis

10 μg/mL Hemagglutinin was precomplexed with strepmab and goat-anti-human Alexa Fluor 488 in a 1:0.65:0.325 molar ratio on ice for 30 min. The precomplexed proteins were incubated with 50,000 HCK or HCK-B3GNT2 cells for 30 min at 4°C in the dark. Cells were washed with FACS buffer (PBS containing 1% FCS and 2 mM EDTA) and followed by the centrifuge at 300 rcf for 5 min. The cells were fixed with 1% Paraformaldehyde for 10 min at 4°C in the dark. After being washed with FACS buffer, the cells were resuspended in 100 μL of FACS buffer. Flow cytometry was performed using BD FACSCanto II or the CytoFLEX LX (Beckman Coulter). Data was analyzed using FlowJo software. All cells, single cells, and live cells were gated. Mean fluorescence values for triplicates were averaged, and standard deviations were calculated.

### Molecular dynamics simulations

Monomeric models of HAs from A/SG/16 and its mutants Y159N/T160I and Y159N/T160I/Y195F were generated with AlphaFold2^45^ version 2.3.2 using the following options: *--models-to-relax=all*, *--enable-gpu-relax*, *--db-preset=reduced_dbs*, and *--model-preset=monomer*. For each mutant and wild type, thirty structures were generated for models 3, 4, and 5 (90 structure predictions per sequence). Predicted structures were ranked according to the pLDDT score, and the highest-scoring structure for each sequence was used as the starting point for extensive conventional molecular dynamics (MD) simulations. Models were cropped to include residues 49 to 278 and have a pLDDT score > 95, indicating high prediction confidence.

N158 of the A/SG/16 model was glycosylated with a biantennary symmetric, not core-fucosylated, eptasaccharide, using the GLYCAM Carbohydrate Builder^46^. For A/SG/16 and its Y159N/T160I/Y195F mutant, initial geometries of the complex with **D** were prepared, generating the coordinates of **D** using the GLYCAM Carbohydrate Builder^46^ followed by manual overlay onto the AlphaFold2 protein model.

MD simulations were carried out with AMBER 24^47^ using the ff14SB force field for the proteins^48^, the GLYCAM-06j force field for glycans^49^, and the TIP3P force field for water^50^. Unbound and bound models were immersed in a water box with a 10 Å buffer of water molecules and neutralized by adding explicit Cl^-^ counterions (Li-Merz 12-6 normal usage set)^51^. A two-stage geometry optimization approach was implemented. The first stage minimizes only the positions of solvent molecules and ions, and the second stage is an unrestrained minimization of all the atoms in the simulation cell. The systems were then heated by incrementing the temperature from 0 to 300 K under a constant pressure of 1 atm and periodic boundary conditions. Harmonic restraints of 10 kcal mol^-1^ Å^-2^ were applied to the solute, and the Andersen temperature coupling scheme was used to control and equalize the temperature^52,53^. The time step was kept at 1 fs during the heating stage, allowing potential inhomogeneities to self-adjust. Once equilibrated, the system was subjected to a 2 ns constant volume molecular dynamics simulation at 300 K using the SHAKE algorithm and a 2 fs time step^54^. Long-range electrostatic effects were modeled using the particle mesh Ewald method^55^. A cutoff of 8 Å was applied to Lennard-Jones interactions. Production was run with the same setup as three independent 400 ns replicas, for a global simulation time of 1.2 μs per system. Analysis was performed with cpptraj^56^.

## Acknowledgments

RL is supported by a CSC fellowship. F.P. is supported by MCIN/AEI grants RYC2022-036457-I and EUR2023-14346. Research reported in this publication was supported by the National Institute of Allergy and Infectious Diseases of the National Institutes of Health under Award Number R01 AI165692 (to G.J.B). The content is solely the responsibility of the authors and does not necessarily represent the official views of the National Institutes of Health. This research was funded by the European Research Council (ERC-2023-STG, project number 101117639-Glyco13Cell to L.U.)

