## Supplemental figures for "Epistasis in the receptor binding domain of contemporary H3N2 viruses that reverted to bind sialylated diLacNAc repeats"

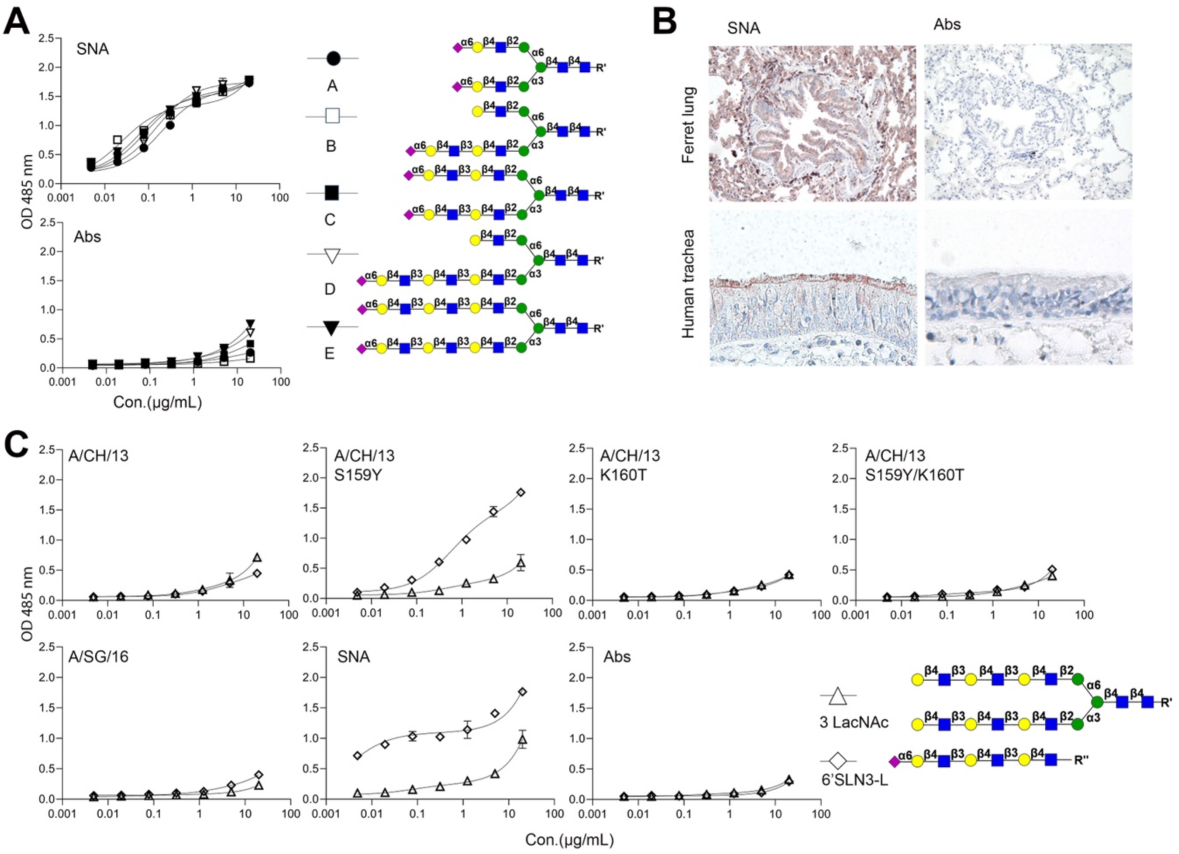


**Figure S1. The correlation between glycan lengths with HA Y159/T160. A.** Bining avidities SNA and antibodies only (Abs) were measured by ELISA. **B.** Tissue staining of SNA and Abs to ferret lung and human trachea. **C.** Binding avidities to 6' SLN3-L and 3LacNac without sialic acid for A/CH/13 WT and mutants, A/SG/16, SNA, and Abs.

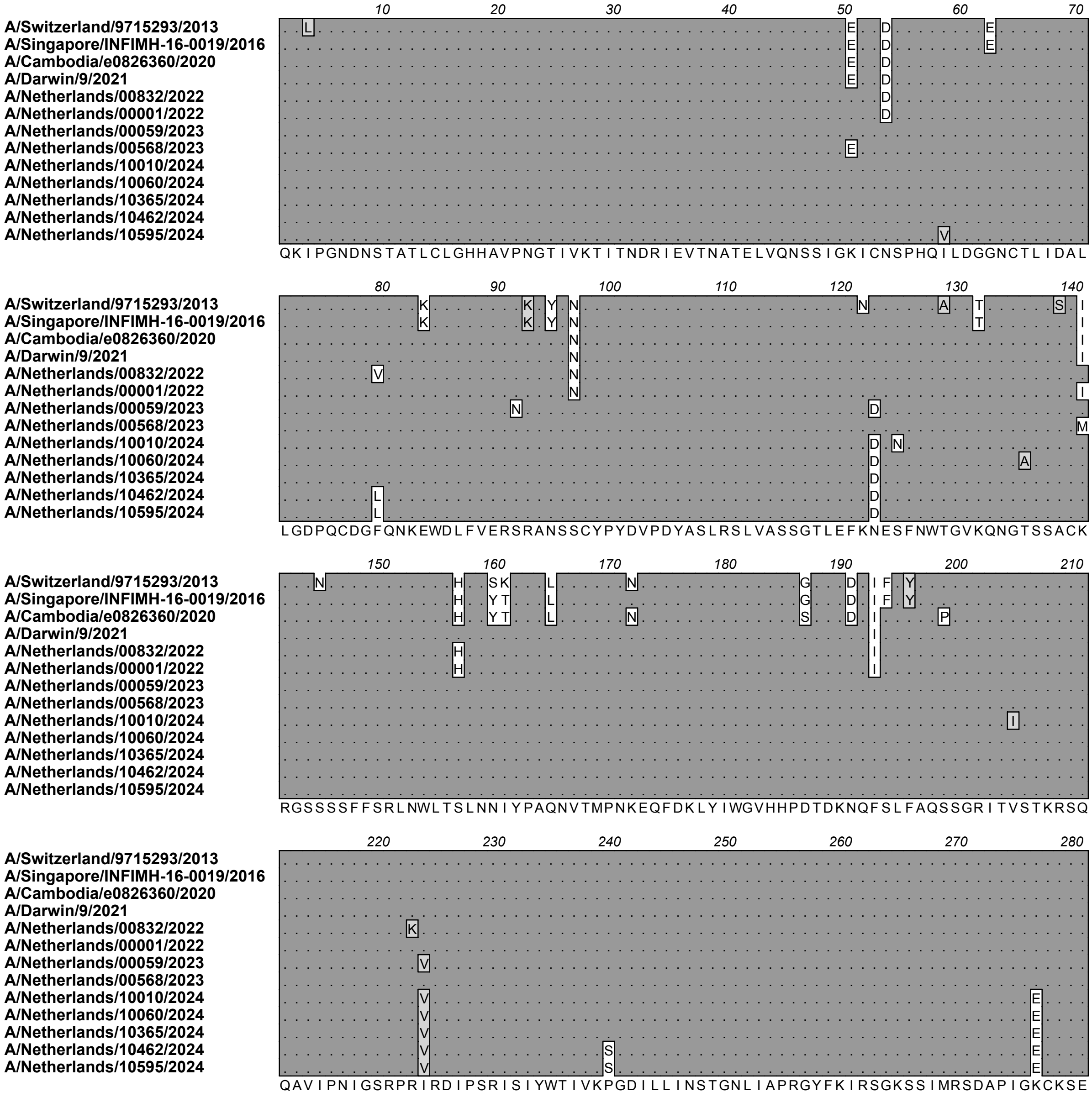


**Figure S2. amino acid aligment of HA1 (1-280) comparing vaccine strains with recent circulating H3N2 viruses.**

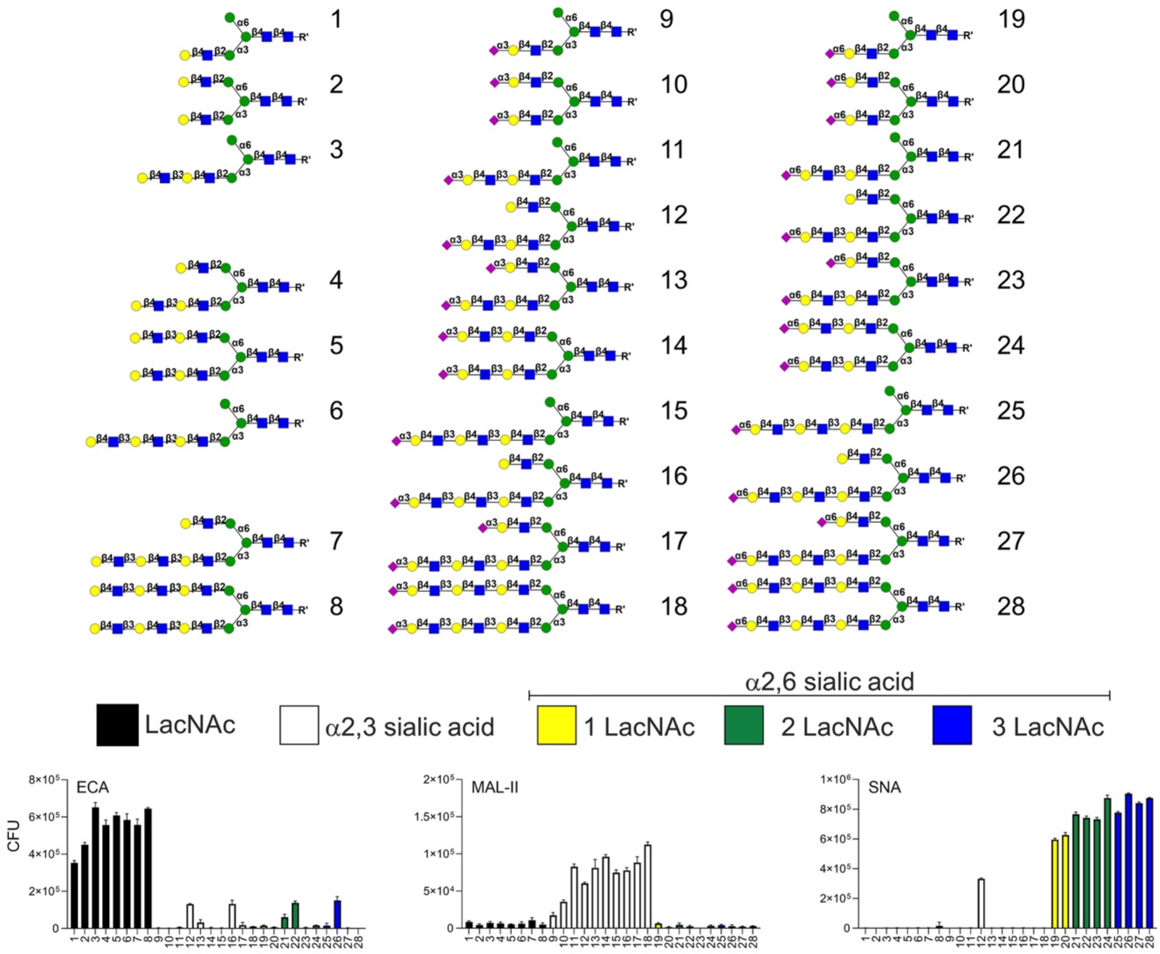


**Figure S3. Population of the the glycan array and quality control thereof**. ECA recognizes terminal galactose (**#1-8, 12 ,16, 22** and **26**). MAL-II recognizes the glycans terminating with α2,3 sialic acid **(#9-18**). SNA recognizes the glycans terminating with α2,6 sialic acid (**#19-28**).

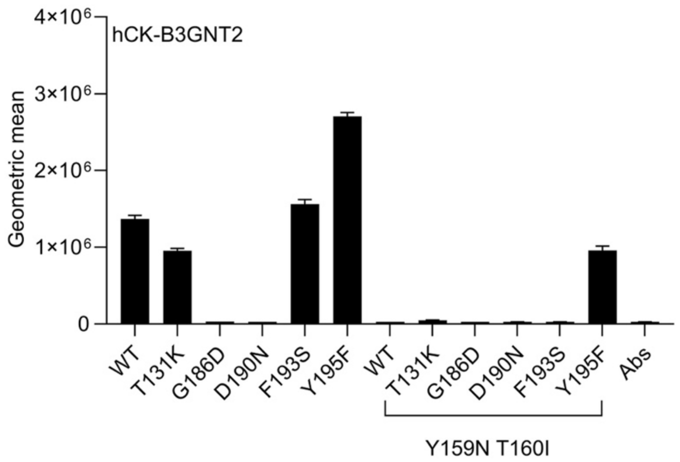


**Figure S4. Flow cytometry analysis of binding to hCK-BGNT2 cells.**

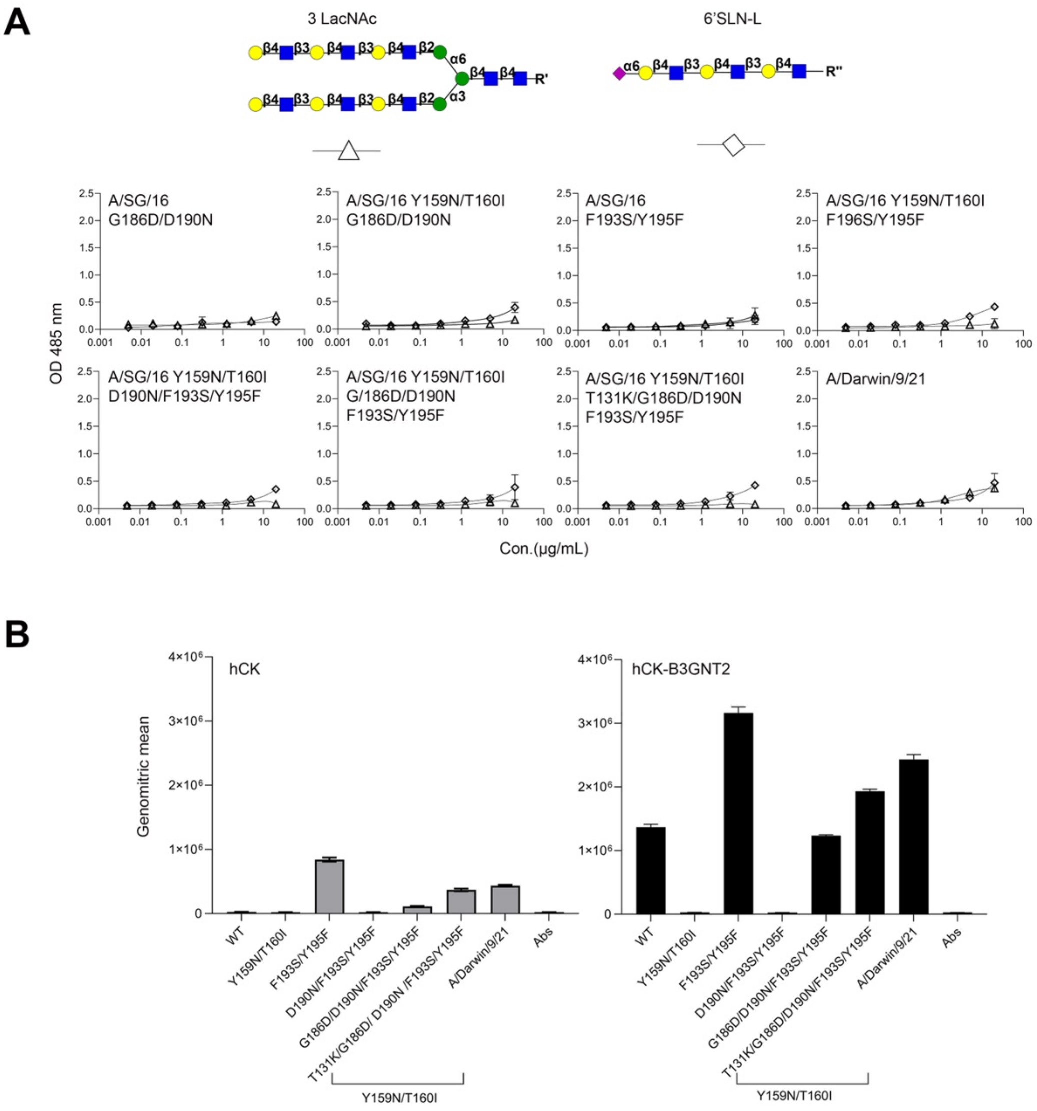


**Figure. S5. Binding activity of different mutant based on the background of A/SG/16 or A/SG16 Y159N/T160I. A**. Binding avidities to 6' SLN3-L and 3LacNAc without sialic acid. **B**. Flow cytometry analysis of binding to CK and hCK-BGNT2 cells.

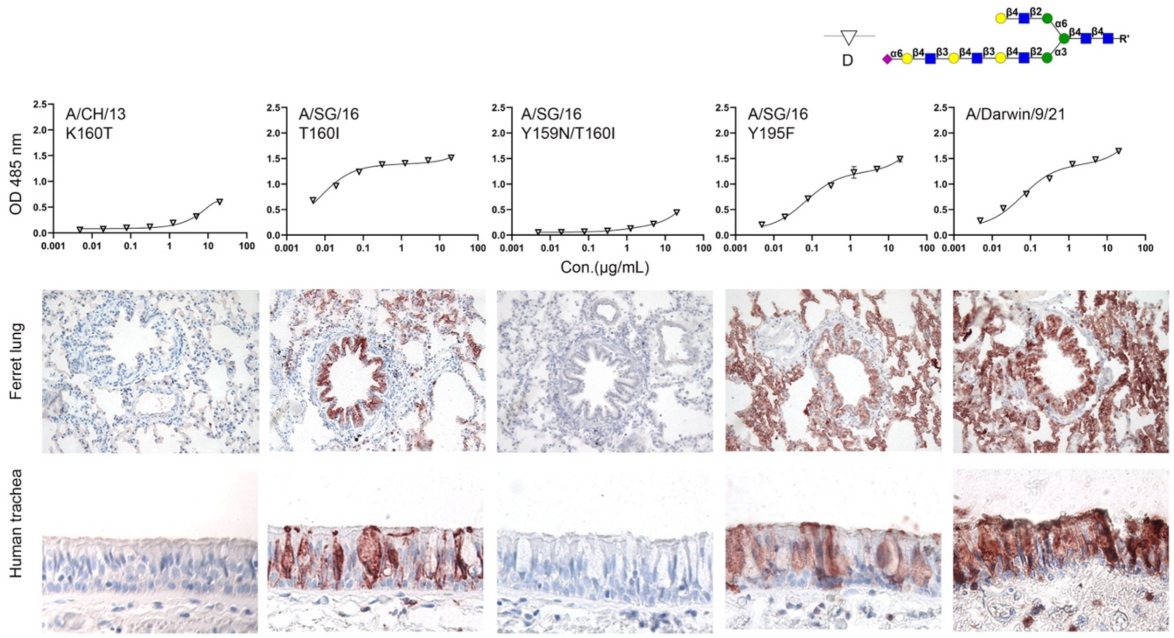


**Fig S6. The relationship between binding to asymmetrical N-glycan with a α2,6 sialylated triLacNAc (compound D) and binding to human and ferret respiratory tissues.** ELISA result (up) *vs* Tissue staining (down).
